# Steady-state, therapeutic, and helminth-induced IL-4 compromise protective CD8 T cell bystander activation

**DOI:** 10.1101/2024.06.10.598293

**Authors:** Nicholas J Maurice, Talia S Dalzell, Nicholas N Jarjour, Taylor A DePauw, Stephen C Jameson

**Affiliations:** Center for Immunology, University of Minnesota Medical School, Minneapolis, MN; Department of Laboratory Medicine and Pathology, University of Minnesota Medical School, Minneapolis, MN

## Abstract

Memory CD8 T cells (T_mem_) can be activated into innate-like killers by cytokines like IL-12, IL-15, and/or IL-18; but mechanisms regulating this phenomenon (termed bystander activation) are not fully resolved. We found strain-intrinsic deficiencies in bystander activation using specific pathogen-free mice, whereby basal IL-4 signals antagonize IL-18 sensing. We show that therapeutic and helminth-induced IL-4 impairs protective bystander-mediated responses against pathogens. However, this IL-4/IL-18 axis does not completely abolish bystander activation but rather tunes the expression of direct versus indirect mediators of cytotoxicity (granzymes and interferon-γ, respectively). We show that antigen-experience overrides strain-specific deficiencies in bystander activation, leading to uniform IL-18 receptor expression and enhanced capacity for bystander activation/cytotoxicity. Our data highlight that bystander activation is not a binary process but tuned/deregulated by other cytokines that are elevated by contemporaneous infections. Further, our findings underscore the importance of antigen-experienced T_mem_ to dissect the contributions of bystander T_mem_ in health and disease.

## Introduction

CD8 memory T cells (T_mem_) are generated to provide rapid protection when their cognate antigen (Ag) is reencountered^1^; however, CD8 T_mem_ are also capable of innate-like, Ag-independent functions when activated by pro-inflammatory cytokines^2–5^. The cellular consequences of this phenomenon, termed bystander activation, include proliferation^6^ and cytotoxicity^7–12^. The cytokines IL-12, IL-15, and/or IL-18 potently elicit bystander-mediated cytotoxicity^2,8,13,14^. Once activated, bystander CD8 T_mem_ indirectly kill by secreting interferon-γ (IFN-γ)^7,8,11,15^ and directly kill by NKG2D-dependent recognition of target cells expressing NKG2D ligands (a family of stress induced proteins)^9,10,16–19^.

However, these effector functions of activated bystander T_mem_ are a double-edged sword^3–5^. Bystander-mediated killing promotes pathogen clearance during acute infections^9,15,16,20,21^. But during sustained/dysregulated inflammation, these responses beget immunopathologies^10,11,17–19,22^. Data thus far suggest the entire pool of CD8 T_mem_, including conventional T_mem_^10,12,17,18,23,24^, tissue-resident T_mem_^15,20^, and Ag-inexperienced “virtual memory” T cells (T_VM_)^16,21,22^, are capable of bystander activation and contributing to innate-like immune responses. Thus, there is significant interest in understanding what specifically predisposes bystander CD8 T_mem_ for enhanced cytotoxicity and what could be used to modulate functionality.

Conventional T cell responses are tightly regulated. Chronic T cell receptor (TCR)-activation leads CD8 T_mem_ to express molecules like TOX and PD-1, which render the cell refractory to additional TCR stimulation in a cell-intrinsic manner^25–29^. Although cytokine-mediated activation transiently increases the expression of both TOX and PD-1 in bystander CD8 T_mem_, these cells remain functional, expressing high levels of IFN-γ and cytotoxic granules like granzyme B^24^. Despite indifference to conventional means of attenuation, bystander activation can be influenced by cell-extrinsic factors. Systematic analysis of cytokine combinations showed IL-4, IL-10, and/or TGFb impair IFN-γ expression during in vitro bystander activation with IL-12, IL-15, and IL-18^14^. Indeed, IL-10 plays a role in regulating bystander activation, as IL-10–deficient animals develop Lyme arthritis that is exacerbated by activated bystander CD8 T_mem_^11^. Nevertheless, the biologic significance of signals like IL-4, especially during acute infection, remain unclear.

By surveying multiple inbred mouse strains in specific pathogen-free (SPF) conditions, we find strain-intrinsic propensities for CD8 T_mem_ bystander activation. We demonstrate that strain-specific deficiencies in bystander activation are partially caused by basal IL-4 signaling, which antagonizes the expression of the cytokine receptor IL-18, which is necessary for aspects of bystander activation in CD8 T_mem_. Using infection models, we show that even in strains predisposed for bystander-mediated cytotoxicity, exogenous or helminth-induced IL-4 can mute protective bystander responses like IFN-γ secretion. Though we confirm the in vivo role of IL-4 in limiting bystander-derived IFN-γ, we show that IL-4 does not fully abrogate bystander activation but rather tunes the bystander CD8 T_mem_ response by re-shaping cytokine receptor expression. We dissect these mechanisms in Ag-experienced animals and demonstrate that strain-intrinsic propensities in bystander activation are overcome by conventional memory differentiation, which enforces the expression of cytokine receptors necessary for bystander activation. We discuss the relevance of IL-4-mediated tuning of bystander responses in the context of health, disease, therapies, and contemporaneous infection as well as the limitations of SPF models for understanding the full contribution of activated bystander CD8 T_mem_ in beneficial and deleterious immune responses.

## Results

### Inbred mouse strains differentially respond to bystander-activating cytokines

We first asked whether strain-specific propensities for bystander activation exist in CD8 T_mem_ from widely-used, commercially-available, inbred mouse strains (C57B6 and BALB/c) as well as their F1 crosses (CB6F1). To minimize T cell-extrinsic signals that could impact activation^30^, we isolated CD8 T cells from secondary lymphoid organs, which we stimulated with IL-12, IL-15, and IL-18 in combination (IL-12/15/18) (**Figure 1A**). Though memory phenotype (i.e., CD44^hi^) CD8 T cells respond to IL-12/15/18 by upregulating effector molecules, IFN-γ and granzyme B, and the activation biomarker, CD69, we found the magnitude of these responses to be strain-dependent, ranging from highest in C57BL/6 to lowest in BALB/c, with the cross of the two having intermediate expression of these markers (**Figure 1B, C; Supplementary figure 1A**).

**Figure 1.**
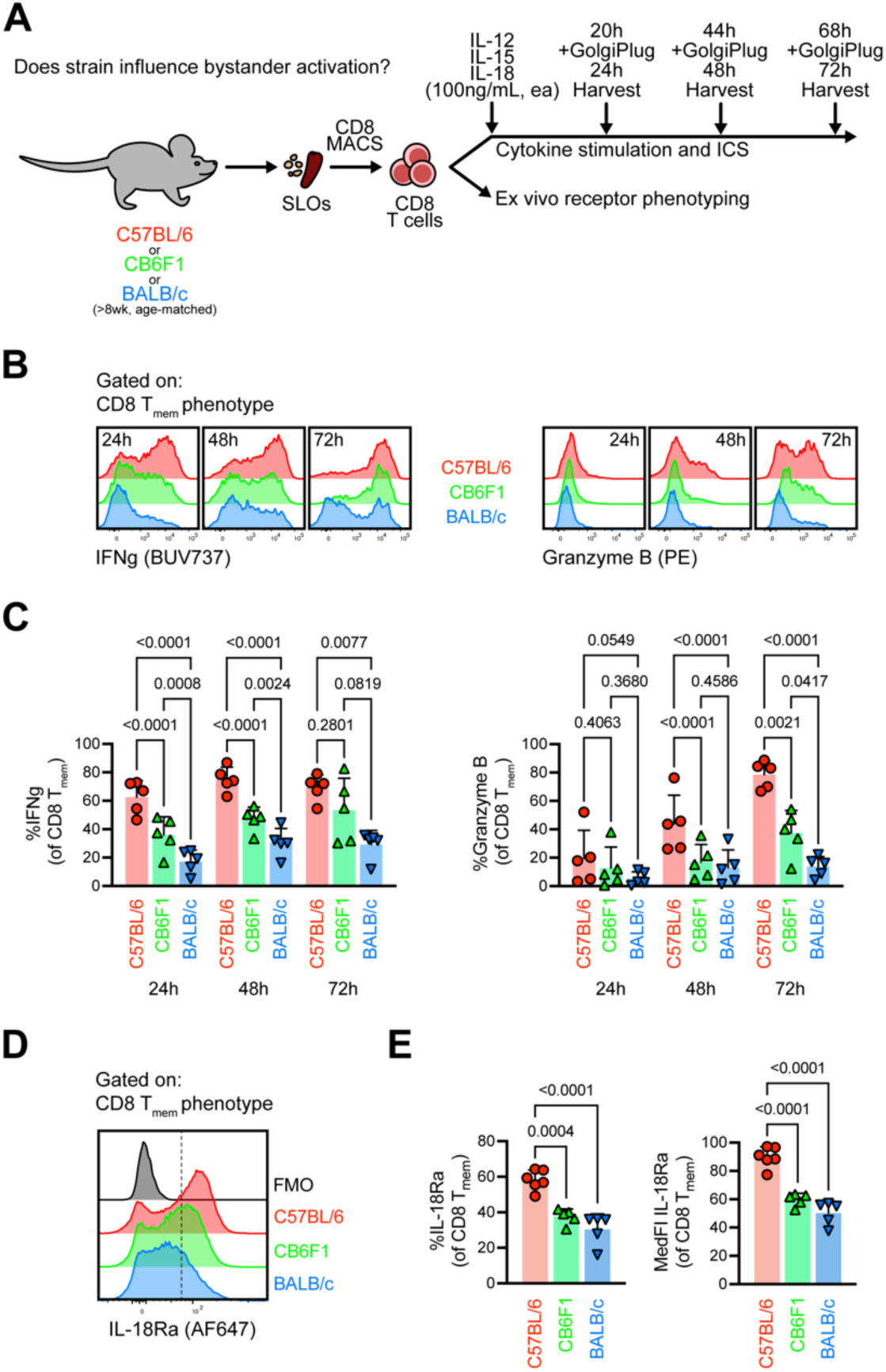
Bystander-activating cytokines differentially activate CD8 T_mem_ from SPF laboratory mice. **A** Experimental layout for CD8 T cell stimulations and ex vivo receptor phenotyping, in which we isolated bulk CD8 T cells using MACS before stimulating with the bystander-activating cytokines, IL-12, IL-15, and IL-18 in combination (100ng/mL, ea). We added GolgiPlug 4 hours prior to collecting cells for flow analysis. **B** and **C** IFN-γ and Granzyme B expression in memory phenotype (CD44^hi^) CD8 T cells after stimulation with bystander-activating cytokines. **D** Basal expression of IL-18Ra in CD8 T_mem_ across SPF laboratory animals. Each point depicts a unique animal at a distinct timepoint (n=5 per strain) across 2 technical replicates. Indicated statistical significance was calculated by Kruskal-Wallis tests with Dunn’s multiple comparisons tests.

We considered if this reflected overall strain-specific deficiencies in activation, but this was not the case, as CD8 T_mem_ uniformly upregulate CD69 after TCR agonism, regardless of strain (**Supplementary figure 1A**). Although mice were housed in specific pathogen-free (SPF) conditions, we excluded the role of strain-intrinsic microbiota on CD8 T_mem_ responses by co-housing animals for ≥3 weeks before conducting stimulation assays (**Supplementary figure 1B**). We found that despite SPF co-housing, strain-specific CD8 T_mem_ responses (i.e., IFN-γ and granzyme B upregulation) in response to IL-12/15/18 remained intact, highlighting probable genetic determinants of bystander activation. Given the abundance of Ag-inexperienced “virtual memory” (T_VM_) cells in SPF animals, we investigated if T_VM_ demonstrate strain-specific deficiencies in bystander activation. We identified T_VM_ by low CD49d and observed T_VM_ from BALB/c animals were indeed lowest for IFN-γ and CD69 expression after in vitro exposure to IL-12/15/18 (**Supplementary figure 1C**).

Since cytokine sensing is requisite for bystander activation, we interrogated the ex vivo expression of cytokine receptor components (**Figure 1A**). We found IL-18Ra expression mirrored the magnitude of responses to IL-12/15/18, wherein its expression was highest in C57BL/6- and lowest in BALB/c-derived CD8 T_mem_ and T_VM_ (**Figure 1D, Supplementary figure 1D**). Thus, IL-18Ra expression patterns were consistent with a role for altered IL-18 sensing driving strain-specific propensities for bystander activation. We hypothesized that this was due in part to IL-4 for dual reasons. First, BALB/c mice demonstrate higher levels of steady-state IL-4 expression^31^; second, exogenous IL-4 was shown to limit IFN-γ expression in BALB/c CD8 T_mem_ that were bystander activated in vitro^14^.

### Steady-state IL-4 signaling partially underlies BALB/c deficiencies in bystander activation

Given the poor responses of BALB/c CD8 T_mem_ stimulated by IL-12/15/18 (**Figure 1B**) and their heightened basal levels of IL-4^31^, we asked if restricting IL-4 signals in BALB/c mice improves the ability to become bystander activated. We utilized germline-deficient IL-4Ra BALB/c animals, which lack the receptor subunit required for IL-4 signaling, and found that steady state IL-18Ra expression was elevated in CD8 T_mem_ from *Il4ra^-/-^* BALB/c animals (**Figure 2A**). Given the higher IL-18Ra expression on *Il4ra^-/-^* BALB/c CD8 T_mem_, we asked if this rendered cells more sensitive to IL-12/15/18-mediated activation. We stimulated CD8 T cells from littermates from a cross from *Il4ra^+/-^*BALB/c mice before interrogating IFN-γ expression (**Figure 2B**). As expected, *Il4ra^-/-^*cells were insensitive to IL-4-mediated inhibition of bystander activation in vitro (**Figure 2C**). But we also found that even in the absence of IL-4 in culture conditions, *Il4ra^-/-^* BALB/c CD8 T_mem_ have enhanced IFN-γ expression in response to IL-12/15/18 in comparison to *Il4ra^+/+^*and *Il4ra^+/-^* littermate controls at the 24-hour timepoint (**Figure 2C**). This advantage in *Il4ra^-/-^*BALB/c CD8 T_mem_ waned at later timepoints (**Supplementary figure 2**), possibly because cells capable of IL-4 sensing have been removed from the tonic IL-4 signals that occur in vivo. Despite the role of basal IL-4 signaling in poor BALB/c bystander activation, other factors also likely regulate bystander activation, as CD8 T_mem_ from C57BL/6 mice still demonstrate greater IFN-γ responses (**Figure 2C**). Though we show that IL-4 plays a role in limiting BALB/c bystander activation at steady state, we wondered if exogenous IL-4 would impair IFN-γ and IL-18Ra expression in strains prone to bystander activation.

**Figure 2.**
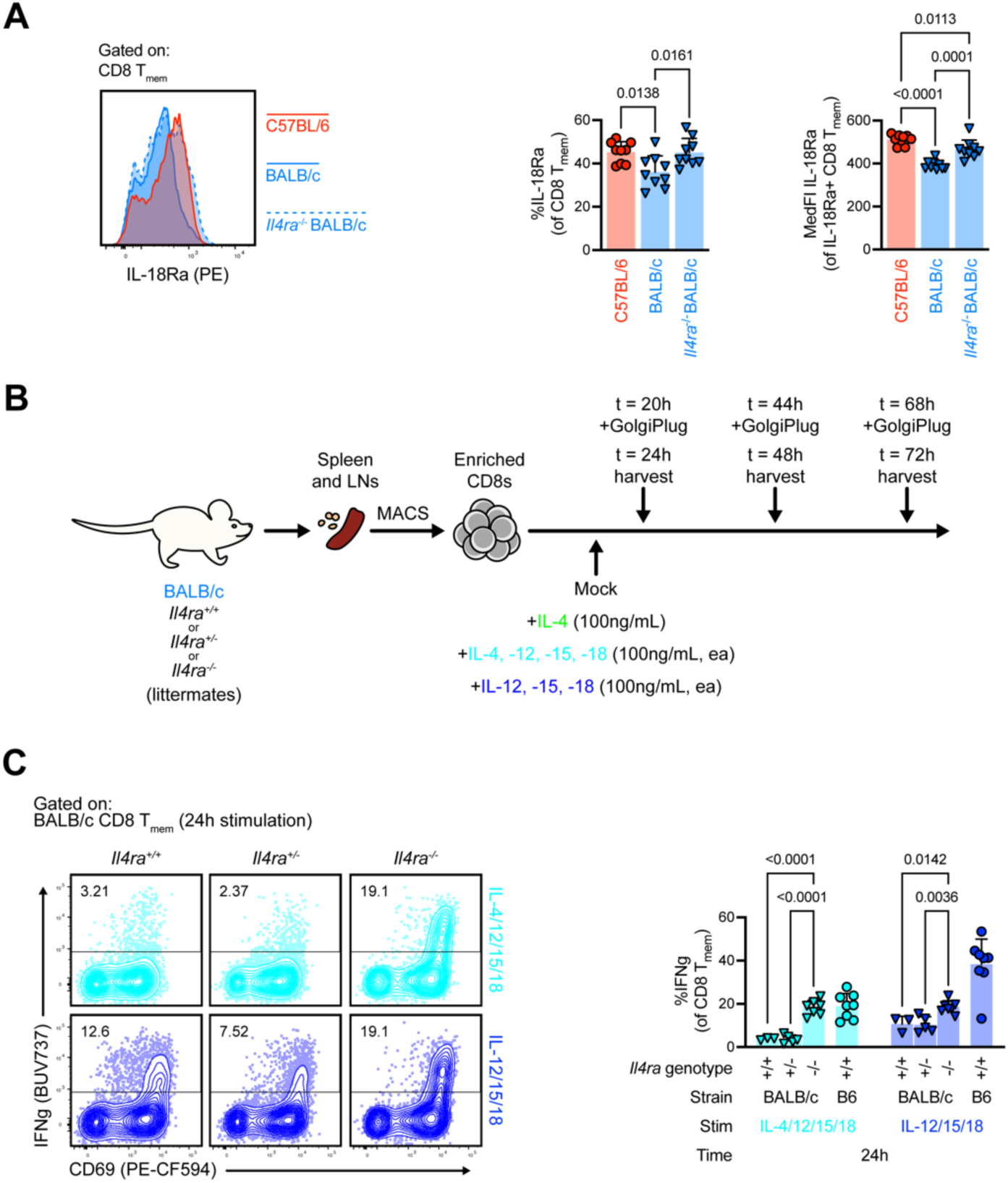
Disrupting steady-state IL-4 signaling in BALB/c CD8 T_mem_ partially restores response to bystander-activating cytokines. **A** Steady-state IL-18Ra expression in blood CD8 T_mem_ from C57BL/6, wildtype (*Il4ra^+/+^*) BALB/c, or *Il4ra^-/-^* BALB/c animals. **B** Experimental diagram outlining CD8 T cell isolation from *Il4ra* wildtype, heterozygous, and knockout BALB/c mice and subsequent stimulation with bystander-activating cytokines (IL-12/15/18, 100ng/mL, ea.) with or without IL-4 (100ng/mL). **C** IFN-γ expression in CD8 T_mem_ from BALB/c *Il4ra* variants after 24h of stimulation with IL-4/12/15/18 and IL-12/15/18. **D** Experimental diagram for testing in vivo bystander activation and bystander-mediated pathogen clearance. We depleted NK cells from BALB/c mice using anti-asialo GM1 antibodies before infecting mice with 1,000 CFU wildtype *L. monocytogenes*. We injected animals with fluorophore-conjugated anti-CD8b 3 min prior to harvest to infer T cell location, and collected spleen and liver tissue to test cellular phenotypes and bacterial burden. **E** Ex vivo IFN-γ and IL-18Ra expression in CD8 T_mem_. **F** *L. monocytogenes* CFU within spleens and livers. Each symbol in **A** depicts and individual animal (n=9 per strain), each symbol in **C** depicts cells from an individual animal in a unique stimulation condition (n=3–8 per strain/geneotype) across 2 technical replicates. Indicated statistical significance was calculated by ordinary one-way ANOVA with **A** Tukey’s or **C** Holm-Šídák multiple comparisons tests.

### IL-4 limits IL-18Ra expression and IFN-γ elicited by bystander-activating cytokines

We sought to test the role of IL-4 on bystander activation in C57BL/6 animals using CD8 T_mem_ with defined antigen specificity. To develop a defined CD8 T_mem_ population, we adoptively transferred naïve, congenically-distinct (i.e., *Cd45*^.1/.1^) OT-I CD8 T cells, which express a transgenic TCR that recognizes the SIINFEKL peptide of chicken-egg ovalbumin (OVA), into background-matched C57BL/6 recipients (**Figure 3A**). We infected recipients with an OVA-expressing vesicular stomatitis virus construct (VSV-OVA), to provide the signals needed for memory differentiation. We used mice ≥60 days post VSV-OVA infection (referred to as OT-I memory mice) for subsequent stimulation assays to test the role of IL-4 on IL-12/15/18-mediated activation (**Figure 3A**). We found that IL-4 signaling in tandem with IL-12/15/18 reduced IFN-γ expression in OT-I T_mem_, which was most prominent at 24 hours (**Figure 3B**). This IL-4-mediated phenomenon was mirrored in CD8 T_mem_ from SPF animals, regardless of strain (**Supplementary figure 3A, B**), suggesting that the ability of IL-4 to limit responses to bystander-activating cytokines is not a strain-specific phenomenon. Despite this, we found that the degree to which IL-4 impaired IL-12/15/18-induced IFN-γ was greatest in BALB/c CD8 T_mem_, suggesting that both IL-4 levels as well as sensitivity to IL-4 may confer limited bystander activation in CD8 T_mem_ from BALB/c animals (**Supplementary figure 3C**).

**Figure 3.**
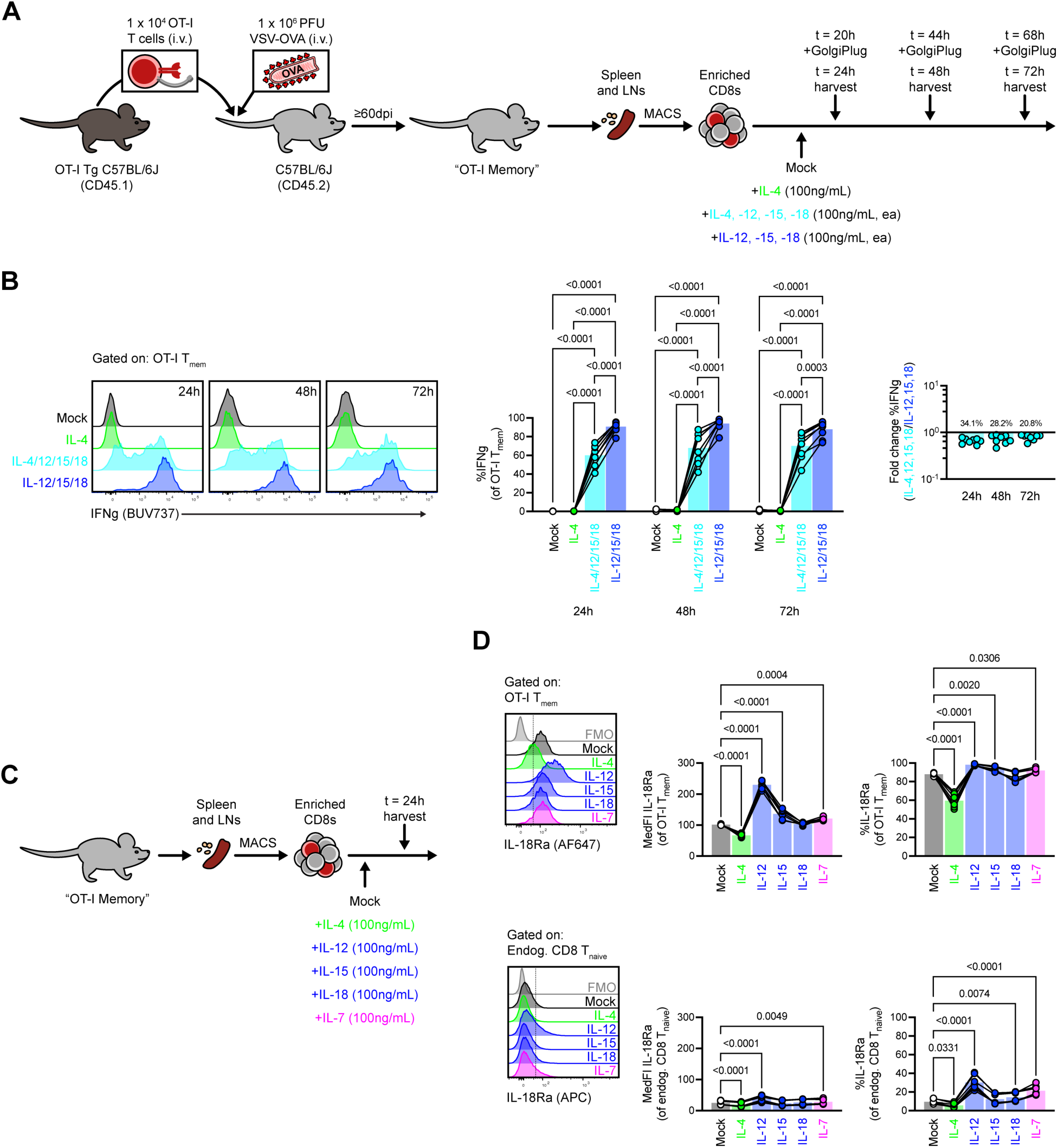
IL-4 limits IL-18Ra expression and IFN-γ expression elicited by bystander-activating cytokines. **A** Experimental diagram outlining OT-I T_mem_ generation and subsequent CD8 T cell isolation and in vitro activation with bystander-activating cytokines (IL-12/15/18, 100ng/mL, ea.) in the presence or absence of IL-4 (100ng/mL). **B** IFN-γ expression in OT-I T_mem_ after cytokine stimulation. **C** Experimental diagram for cytokine stimulation and IL-18Ra phenotyping, in which we stimulated bulk CD8 T_mem_ from OT-I memory mice with single cytokines (100ng/mL) for 24h before flow interrogation. **D** Expression of IL-18Ra in OT-I T_mem_ and endogenous CD8 T_naive_ after 24h stimulation with single cytokines. Each point in **B** and **D** depict a cells from a single animal at a distinct timepoint, which are connected by animal identity (n=7–8 across 3 technical replicates). Indicated statistical significance was calculated by Friedman tests and Dunn’s multiple comparisons tests comparing **B** all data groups to one another or **D** all data groups against mock-treated cells.

Since IL-4 attenuated IFN-γ expression elicited by IL-12/15/18, we asked if this is due to IL-4 signaling antagonizing the expression of other cytokine receptors. We stimulated CD8 T cells from OT-I memory mice with single cytokines, IL-4, IL-12, IL-15, IL-18, as well as IL-7 (a different yc cytokine related to IL-4 and IL-15) and interrogated OT-I T_mem_ expression of cytokine receptors (**Figure 3C**). While OT-I T_mem_ uniformly express IL-18Ra, we found that exposure to IL-4 alone downregulates IL-18Ra (**Figure 3D**), highlighting that IL-4 can alter the sensitivity to bystander-activating cytokines and the subsequent expression of factors like IFN-γ.

### IL-4 complex therapies limit protective functions of bystander CD8 T_mem_

Given that IL-4 could limit IL-18 sensing and subsequent IFN-γ expression during bystander activation across all strains, we asked if perturbing IL-4 sensing affects the consequences of bystander activation in vivo. To do this, we tested the innate-like control of pathogen conferred by bystander-activated CD8 T_mem_ using a *Listeria monocytogenes* infection model, where the early burst of IFN-γ from bystander T cells restricts pathogen^7^. We first tested if exogenous IL-4 impairs bystander activation in mouse strains that we had earlier characterized to have high magnitude bystander responses (e.g., C57BL/6 animals). To precisely determine the contribution of bystander T cells in this model, we depleted natural killer (NK) cells using anti-NK1.1 (clone PK136), as they can also produce IFN-γ in an inflammation-dependent manner^23,32^. We then dosed mice with IL-4 complexes (IL-4c) immediately prior to infection with *L. monocytogenes* (**Figure 4A**). We found the short pulse of IL-4c treatment led to elevated pathogen burden in an NK-independent manner, even at the earliest (24h) time point (**Figure 4B**).

**Figure 4.**
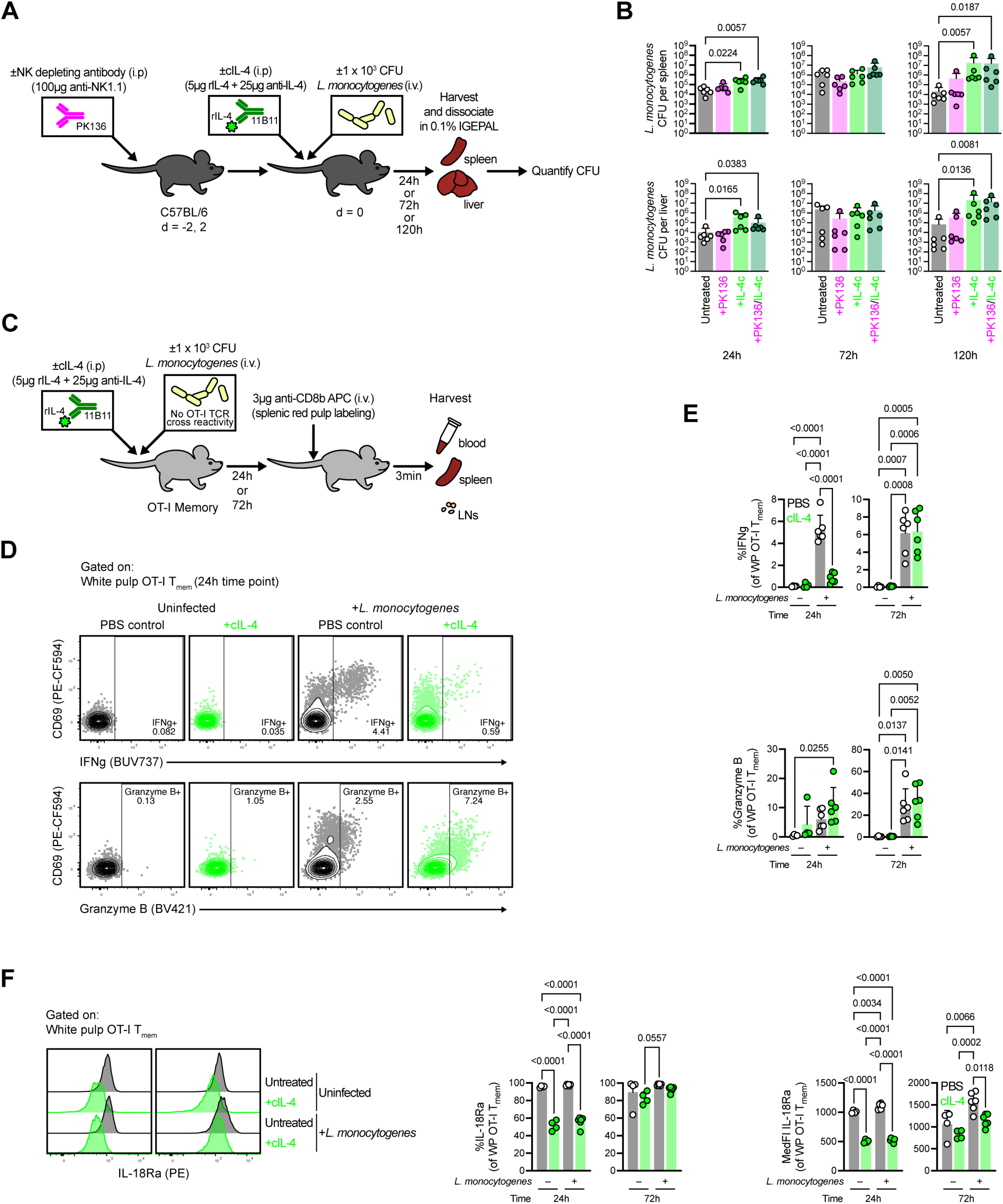
IL-4 complex therapies limit protective functions of bystander CD8 T_mem_. **A** Experiment outline for establishing the effect of IL-4c on *L. monocytogenes* burden during the innate-like immune response, in which we depleted NK cells from C57BL/6 mice with anti-NK1.1 and subsequently infected with 1,000 CFU wildtype *L. monocytogenes*. We then assayed spleen and liver for pathogen at 24, 72, and 120h. **B** *L. monocytogenes* burden in spleen and liver in NK-depleted and/or IL-4c-treated C57BL/6 animals. **C** Experiment outline to test the effect of IL-4c on in vivo bystander activation. We treated OT-I memory mice with IL-4c and infected with *L. monocytogenes*. We injected animals with fluorophore-conjugated anti-CD8b 3 min prior to euthanasia to identify OT-I T_mem_ in the splenic white pulp (WP) and then harvested tissue for flow analysis. **D–F** IFN-γ, granzyme B, and **G** IL-18Ra expression in OT-I T_mem_ at the splenic WP during combined IL-4c treatment and *L. monocytogenes* infection. Each point in **B**, **E**, and **F** represent an individual animal (n=3–6 per timepoint and condition) across 2 technical replicates. Indicated statistical significance was calculated by **B** Kruskal Wallis and Dunn’s multiple comparisons test or **E**, **F** ordinary one-way ANOVA with Tukey’s multiple comparisons tests.

Since IL-4c exacerbated listeriosis in an NK-independent manner, we asked if this correlates with impairment of IFN-γ expression in bystander CD8 T_mem_. We utilized OT-I memory mice, as OT-I T_mem_ lack TCR cross-reactivity with *L. monocytogenes* antigens^9^. We treated mice with IL-4c before infecting with low-dose *L. monocytogenes*, so that early inflammation would be limited to discrete foci of the splenic white pulp (WP), wherein bystander activation occurs^23^ (**Figure 4C**). Prior to harvest, we injected mice intravenously with fluor-conjugated anti-CD8b, so that we could discriminate OT-I T_mem_ within the WP (the site of infection and inflammation), from those in circulation (**Figure 4C**)^23,33,34^. IL-4c treatment reduced IFN-γ expression in bystander OT-I T_mem_ in the WP 24 hours after *L. monocytogenes* infection (**Figure 4D, E**); though these effects were less pronounced at the 72-hour timepoint, mirroring our in vitro stimulation data, in which IL-4-mediated IFN-γ impairment was constrained to earlier timepoints (**Figure 2C**, **Supplementary figure 2B**); though this could also be due to IL-4c clearance. However, counterintuitively, IL-4c treatment led to an upregulation of granzyme B (**Figure 4D, E**). We observed similar patterns in endogenous (i.e., non–OT-I) CD8 T_mem_ and NK cells, wherein IL-4c impaired IFN-γ expression while enhancing granzyme B expression (**Supplementary figure 4A–J**). We asked whether in vivo IL-4c treatment has a similar effect on IL-18Ra expression as we observed in vitro. OT-I T_mem_ from both uninfected and infected animals lost IL-18Ra expression when treated with IL-4c; however, IL-18Ra levels recovered over time (**Figure 4F**), coinciding with the return to normal levels of IFN-γ at later timepoints in IL-4c treated mice (**Figure 4D, E, Supplementary figure 4A– F**).

The IFN-γ secreted by bystander-activated T_mem_ orchestrates the microbicidal functions of antigen-presenting cells (APCs)^35,36^, so we asked if impaired IL-4-mediated impairment of IFN-γ expression in bystander T_mem_ correlates with attenuated APC function. We first determined that IL-4c treatment alone does not lead to a rapid and substantial expansion of APC subsets that could otherwise explain increased susceptibility to *L. monocytogenes* infection (**Supplementary figure 5A**). However, we found that IL-4c treatment in conjunction with *L. monocytogenes* infection led to a decreased frequency of pro-inflammatory Ly-6C^hi^ monocytes (**Supplementary figure 5B**). Ly-6C^hi^ monocytes are critical for host defense during listeriosis^35,36^, so we asked if IL-4c treatment impairs their IFN-γ-dependent antimicrobial functions. We interrogated the expression of iNOS, an IFN-γ-induced molecule which produces radical nitrogen species, and found that it was only expressed in Ly-6C^hi^ monocytes from infected animals (**Supplementary figure 5C**); and a lower proportion of iNOS-expressing cells were present in infected mice that were simultaneously treated with IL-4c (**Supplementary figure 5C**). Upon deeper phenotyping, we found iNOS-positive Ly-6C^hi^ monocytes upregulated other IFN-γ-inducible markers, like CD86 and PD-L1 (**Supplementary figure 5D**). This led us to question if IFN-γ-inducible antimicrobial phenotypes directly correlated with IFN-γ levels in bystander OT-I T_mem_. Indeed, we found frequency of IFN-γ in bystander OT-I T_mem_ to be an accurate predictor of Ly-6C^hi^ monocyte expression of iNOS, co-stimulatory molecules (CD86 and CD80), and antigen-presentation molecules (MHC-II, I-A^b^) (**Supplementary figure 5E**). Together, these data suggest IL-4c modulation of IFN-γ expression in bystander CD8 T_mem_ can have profound downstream consequences on other immune cells and subsequent pathogen control.

### IL-4 enhances granzyme B expression at the expense of IFN-γ during bystander activation

Given trends of granzyme B upregulation in IL-4c-treated animals (**Figure 4D**), we asked whether this was recapitulated in our in vitro stimulations (**Figure 5A**). Indeed, IL-4 enhanced granzyme B expression while concurrently limiting IFN-γ expression in OT-I T_mem_ (**Figure 5B**), suggesting IL-4 may tune rather than broadly inhibit bystander activation. We systematically tested the contribution of each cytokine on bystander activation using OT-I T_mem_. Although individual cytokines poorly induced IFN-γ or granzyme B, combinations of IL-12, IL-15, and/or IL-18 induced expression of these molecules (**Figure 5C**). IL-12 and IL-15 together potently upregulated granzyme B; but this was impaired (while IFN-γ expression was simultaneously enhanced) when IL-18 was added (**Figure 5C**). IL-4, which impaired IL-18 sensing (**Figure 2D**, **Figure 4G**), appears to reverse these IL-18-mediated effects, restoring granzyme B expression at the expense of IFN-γ (**Figure 5C**).

**Figure 5.**
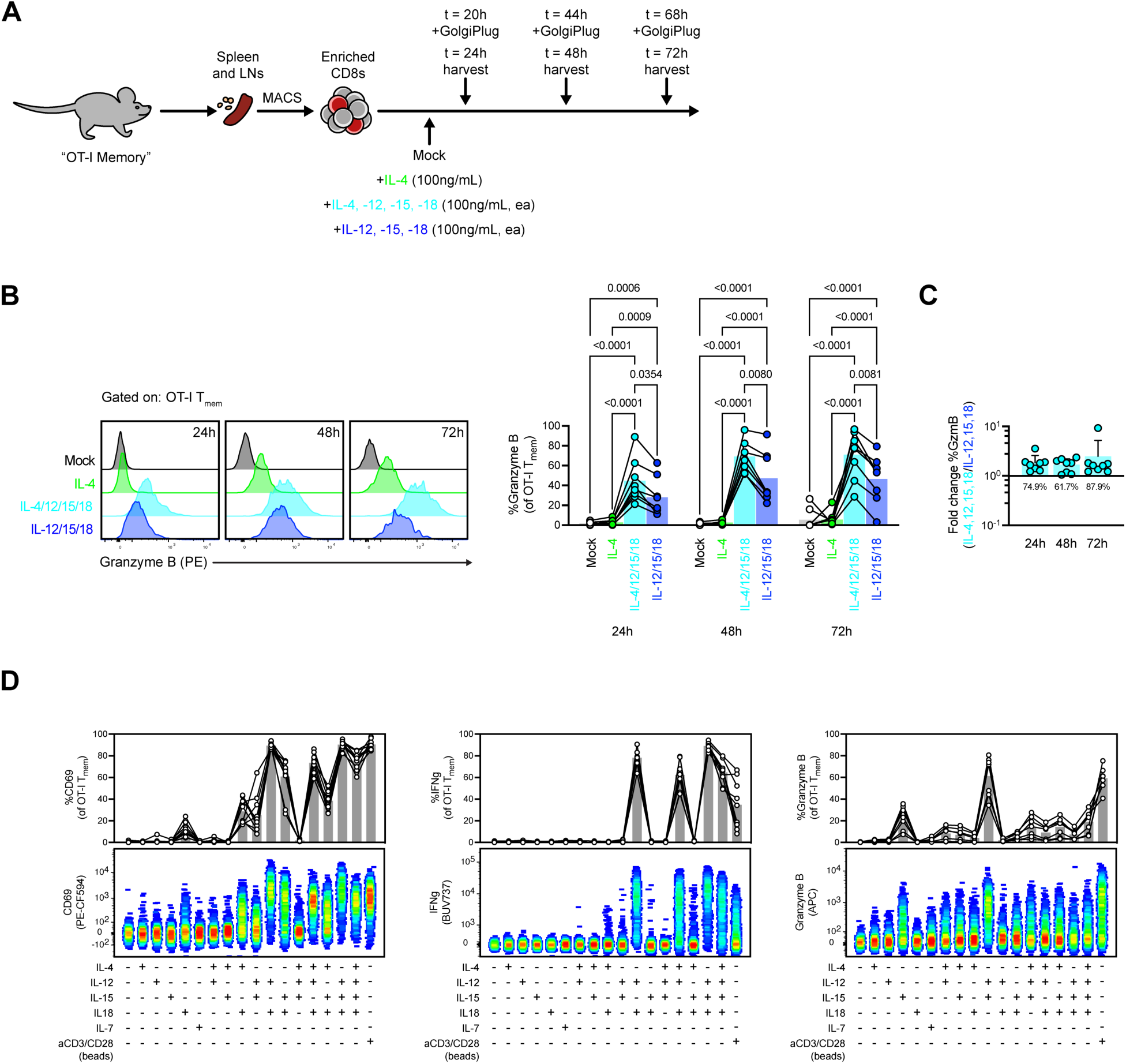
IL-4 enhances granzyme B expression at the expense of IFN-γ during bystander activation. **A** Experimental diagram outlining CD8 T cell isolation from OT-I memory mice and stimulation with IL-4 (100ng/mL) and/or bystander-activating cytokines (IL-12/15/18, 100ng/mL, ea.). **B** Granzyme B expression in OT-I T_mem_ during cytokine stimulation. **C** IL-4-mediated fold change of granzyme B expression elicited in OT-I T_mem_ by bystander-activating cytokines. **D** Expression of CD69, IFN-γ, and granzyme B in OT-I T_mem_ stimulated with various cytokine combinations for 24 hours. Each point in **B**, **C**, and **D** represents an individual timepoint/stimulation connected by animal identity (n=7–9 mice) across 3 technical replicates. Indicated statistical significance was calculated by Friedman tests and Dunn’s multiple comparisons tests.

Since IL-4 can tune bystander T cell functions by regulating IL-18 sensing, we dissected the signaling mechanism responsible for this. IL-4 signaling pathways include JAK/STAT6, MAPK, and PI3-K. We leveraged *Il4ra*^Y500F^ BALB/c animals, which encode an IL-4Ra subunit that cannot signal through the PI3-K pathway^37^, to test its role in tuning bystander activation. Using in vitro stimulations, we found that CD8 T_mem_ from *Il4ra*^Y500F^ BALB/c animals remained sensitive to IL-4-mediated downregulation of IFN-γ and upregulation of Granzyme B expression during IL-12/15/18 stimulation (**Supplementary figure 6A**). Together, these data suggest that IL-4 augments the response to bystander-activating cytokines by tuning sensitivity to IL-18 in a PI3-K independent manner.

Since IL-4c blunted bystander-mediated protection in C57BL/6 mice, we wondered if loss of steady-state IL-4 signaling improves bystander-mediated protection in the BALB/c model. Because BALB/c NK1.1 epitopes fail to react with the PK136 antibody for NK depletions^38^, we depleted NK cells with anti-Asialo GM1. Though efficient at NK depletion, AsGM1 also depleted CD8 T_mem_ (**Supplementary 6B**), given their unexpected AsGM1 expression during homeostasis and activation (**Supplementary 6C, D**). Previous reports showed AsGM1 reactivity with basophils^39^ and stimulated T cells^40,41^, but not on bulk CD8 T_mem_; yet modern studies continue to erroneously leverage AsGM1 as a selective NK-depleting agent. Given the intractability of the BALB/c model in this regard, we returned to the C57BL/6 model to investigate the effects of biologically induced IL-4 on bystander activation.

### Contemporaneous type 2 infections perturb the bystander T cell response to L. monocytogenes

We found that supraphysiologic IL-4 levels can tune the response to bystander-activating cytokines both in vitro and in vivo. While these levels are relevant regarding cytokine-based immunotherapies^42,43^, we wondered if physiologic elevation of IL-4, such as during type 2 infections, can also alter bystander T cell responses. We therefore infected OT-I memory mice with *Heligmosomoides polygyrus*, as this helminth is solely enteric and elicits IL-4 expression^44,45^; we co-infected animals with *L. monocytogenes* at this timepoint (6 days post *H. polygyrus*) and interrogated *L. monocytogenes*-induced bystander activation in the splenic WP 24 hours later (**Figure 6A**). *H. polygyrus* infection alone did not induce CD69 or IFN-γ expression in bystander OT-I T_mem_ in the splenic WP but during *L. monocytogenes* infection, concurrent *H. polygyrus* infection attenuated the expression of CD69 and IFN-γ in bystander OT-I T_mem_ (**Figure 6B**). On the other hand, we observed that *H. polygyrus* infection alone increased granzyme B expression in bystander OT-I T_mem_, an effect which was further enhanced by simultaneous *L. monocytogenes* infection (**Figure 6B**). Since we observed an IL-4:IL-18 axis that respectively tunes granzyme B versus IFN-γ expression during bystander activation, we asked whether type 2 inflammation from *H. polygyrus* infection similarly impaired IL-18 sensing in bystander T cells at sites distal from the gastrointestinal tract. Indeed, we observed blunted IL-18Ra expression in splenic bystander OT-I T cells that was not recovered during *L. monocytogenes* co-infection (**Figure 6C**).

**Figure 6.**
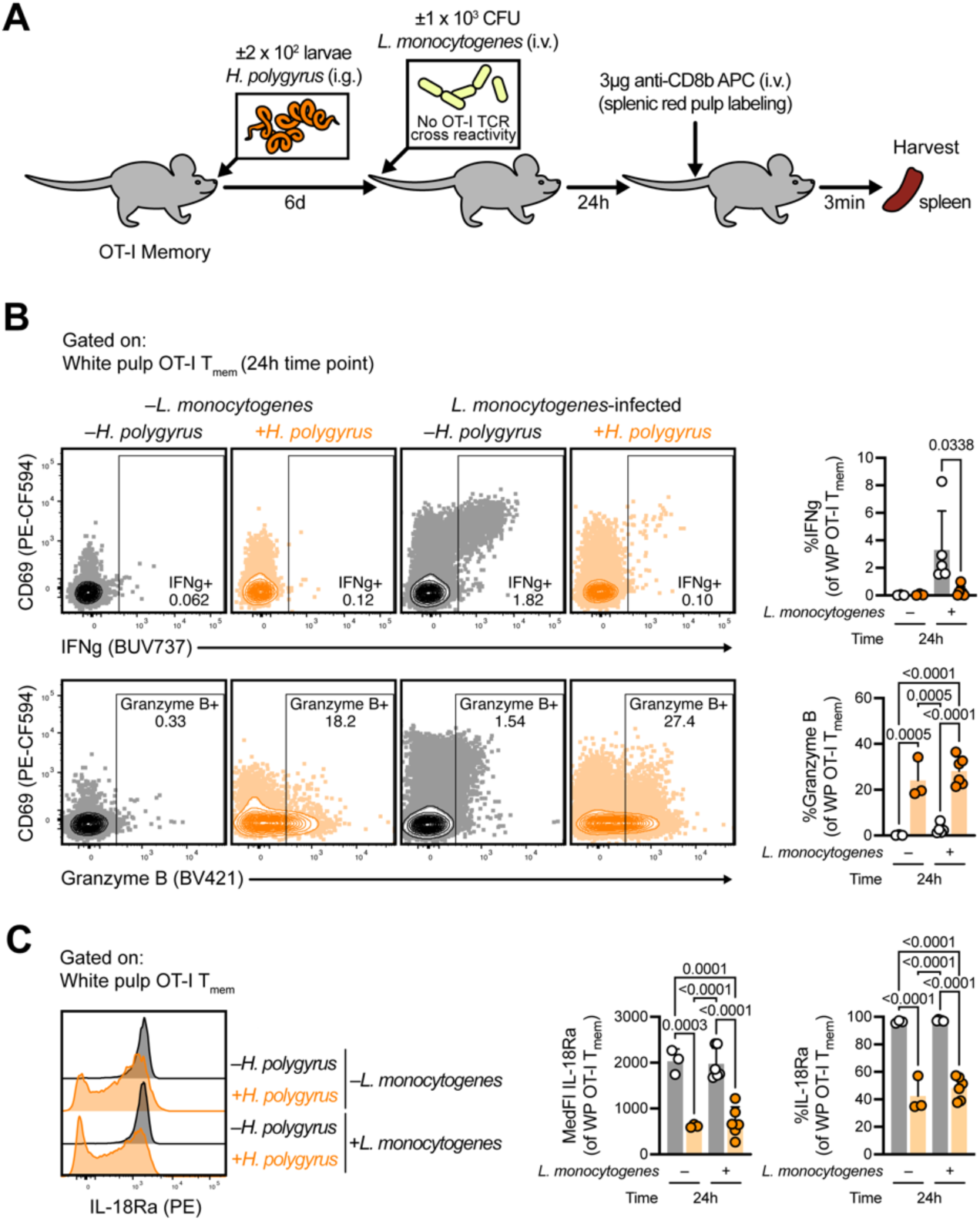
Contemporaneous type 2 infections perturb the bystander T cell response to *L. monocytogenes*. **A** Experimental outline for *H. polygyrus* and *L. monocytogenes* infections. **B** and **C** Ex vivo phenotyping of **B** activation/functional markers and **C** IL-18Ra in bystander OT-I T_mem_ from the splenic white pulp (WP). Each point in **B** and **C** represents an individual timepoint/stimulation connected by animal identity (n=3–6 mice) across 2 technical replicates. Indicated statistical significance was calculated by ordinary one-way ANOVA with Tukey’s multiple comparisons test.

### CD8 T cell priming by viral infection overwrites strain-specific deficiencies in bystander activation

While we determined strain-specific bystander activation to be partially influenced by basal IL-4 signaling, we recognized a limitation was that we were largely employing SPF animals (**Figure 1B**, **Supplementary figure 2A**). Though we employed TCR transgenics, these were limited to the C57BL/6 background, given that the OT-I TCR is restricted to the K^b^ allele of MHC-I. But we observed that OT-I T_mem_ have a functional advantage (i.e., higher IFN-γ expression) in comparison to bulk CD8 T_mem_ (including a large proportion of T_VM_) within the same animal (**Supplementary figure 5C**), suggesting that antigen-experienced cells may be differentially predisposed for bystander activation. To address this, we infected C57BL/6, CB6F1, and BALB/c animals with lymphocytic choriomeningitis virus (LCMV) Armstrong, which causes an acute infection, and then stimulated and interrogated CD8 T cells at a memory timepoint (**Figure 7A**). We observed that LCMV-immune animals still demonstrate strain-specific differences in bystander activation when interrogating bulk, LCMV-nonspecific CD8 T_mem_ (**Figure 7B**). However, when we further analyzed CD8 T_mem_ that expanded in response to LCMV using MHC-I tetramers (e.g., K^d^ GP33–43, K^d^ NP396–404, L^d^ NP118–126) (**Figure 7C**), we found that LCMV-specific CD8 T_mem_ uniformly express IFN-γ in response to in vitro IL-12/15/18, regardless of strain (**Figure 7D**).

**Figure 7.**
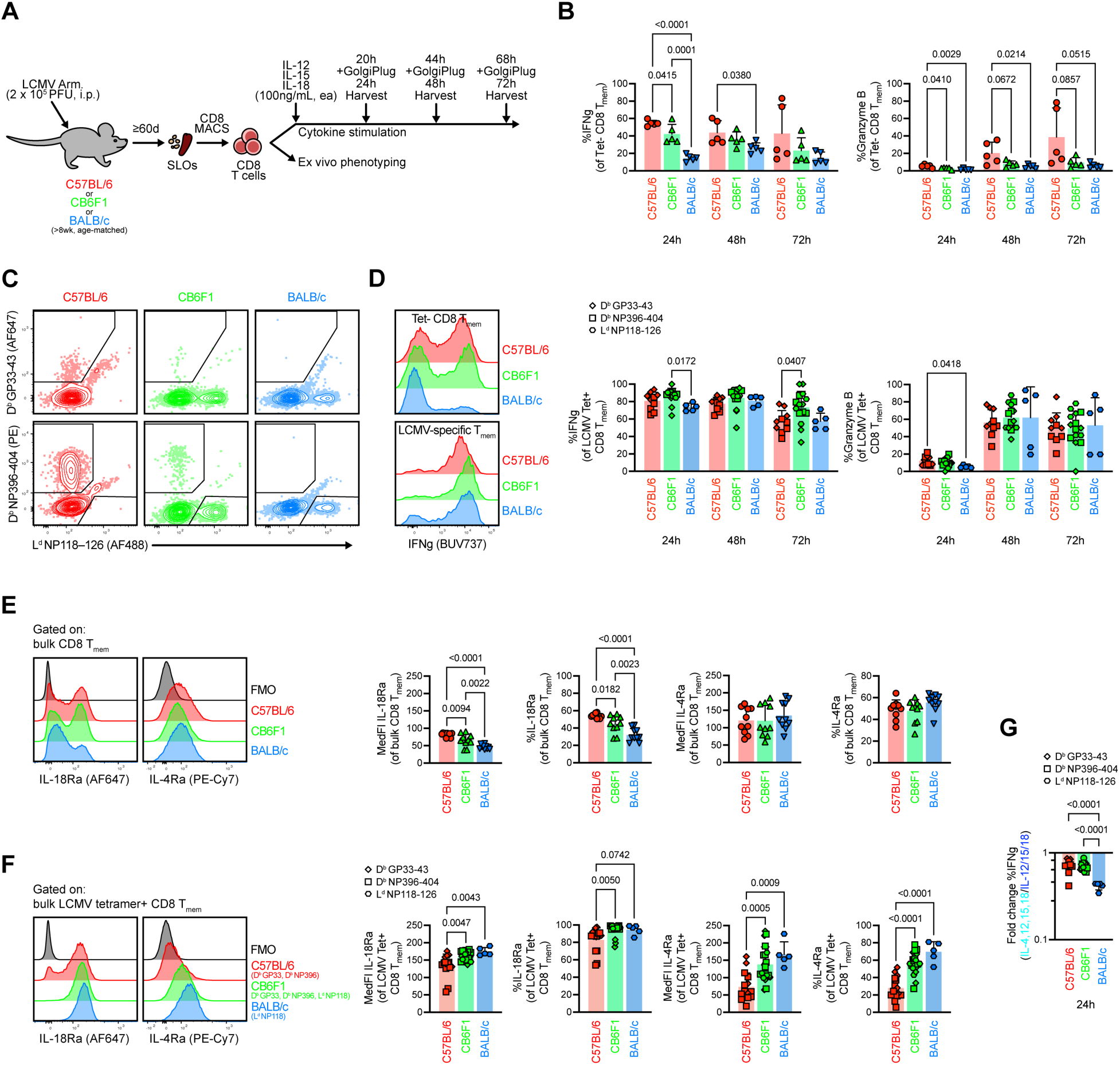
CD8 T cell priming by viral infection overwrites strain-specific deficiencies in IL-18Ra expression and bystander activation. **A** Experiment overview in which we infected C57BL/6, CB6F1, and BALB/c animals with 2 x 10^5^ PFU LCMV Armstrong to develop LCMV-specific CD8 T_mem_. At the memory timepoint, we harvested CD8 T_mem_ for stimulations with bystander-activating cytokines (IL-12/15/18, 100ng/mL, ea.) and ex vivo phenotyping. **B** IFN-γ and granzyme B expression in bulk CD8 T_mem_ during in vitro bystander activation. **C** Representative gating depicting identification of LCMV-specific CD8 T_mem_ with tetramers. **D** IFN-γ and granzyme B expression in LCMV-specific CD8 T_mem_ during in vitro bystander activation. IL-18Ra and IL-4Ra expression in **E** bulk CD8 T_mem_ and **F** LCMV-specific CD8 T_mem_ at homeostasis. **G** IL-4-mediated fold change in IFN-γ expression in LCMV-specific CD8 T_mem_ during stimulation with IL-12/15/18. Each point depicts a unique cell population (i.e., of unique or undefined TCR specificity) from a distinct animal/stimulation (n=5 animals total) across 2 technical replicates. Indicated statistical significance was calculated by ordinary one-way ANOVA with Tukey’s multiple comparisons test.

Since low IL-18Ra expression in CD8 T_mem_ in SPF BALB/c was likely responsible for poor bystander activation, we surmised that LCMV experience reversed this, given the high IFN-γ responses in IL-12/15/18-stimulated LCMV-specific BALB/c CD8 T_mem_ (**Figure 6D**). Indeed, LCMV tetramer-negative CD8 T_mem_ continued to demonstrate differential IL-18Ra expression, which remained lowest in CD8 T_mem_ from BALB/c animals (**Figure 7E**). However, LCMV-specific CD8 T_mem_ expressed high levels of IL-18Ra, independent of animal strain (**Figure 7F**). These data suggest that conventional antigen experience and memory differentiation erases strain-specific nuances in bystander activation by bestowing T cells with uniformly high expression of cytokine receptors. Though strain-specific differences in IL-12/15/18-mediated bystander activation are not found in antigen-experienced CD8 T_mem_, strain-intrinsic sensitivity to additional IL-4 signals remains intact. We found that LCMV-specific CD8 T_mem_ in BALB/c animals expressed higher levels of IL-4Ra (**Figure 7F**) and demonstrated a greater fold reduction in IFN-γ expression when exposed to IL-4 in tandem with IL-12/15/18 (**Figure 7G**).

## Discussion

Bystander T cells contribute to both beneficial and deleterious outcomes when activated into cytotoxic effectors by inflammation; however, the mechanisms that regulate this phenomenon in vivo are not fully resolved. We sought to identify regulators of bystander activation using multiple laboratory strains of mice and interrogating CD8 T_mem_ responses to cytokines responsible for potent bystander activation (IL-12, IL-15, and IL-18). We found that CD8 T_mem_, including T_VM_, from BALB/c mice fail to bystander activate to the same degree as those from C57BL/6 animals in a manner that was partially determined by basal IL-4 signaling that is characteristic of BALB/c mice^31^. Furthermore, increasing IL-4 levels in C57BL/6 animals (which potently bystander activate under normal conditions) impairs IL-18 sensing that is requisite for innate-like IFN-γ secretion, which limits the protective contribution of bystander T_mem_ to *L. monocytogenes* clearance.

Slifka and colleagues had previously shown that IL-4 sensing impairs IFN-γ expression elicited by IL-12/15/18 stimulation in BALB/c animals^14^; however, we provide a mechanistic context for this, by showing that IL-4 stimulation promotes loss of IL-18Ra. Further, we show that IL-4 does not simply abrogate bystander activation by limiting IL-18 signals, but instead biases cells for granzyme expression. Surprisingly, IL-4 from cytokine therapies or contemporaneous helminth infection can skew bystander T cell responses away from IFN-γ expression, which is a critical factor for bystander-mediated protection during *L. monocytogenes* infection. Helminth infection affects a quarter of the world’s population and cytokine therapies are of growing clinical significance; it will be important to consider their effects on protective or detrimental bystander T cell responses as well on the therapeutic tractability of bystander T cells.

This study, to the best of our knowledge, if the first to demonstrate that bystander T cell responses can be impaired by contemporaneous type 2 infections. This contrasts with a study from Rolot and colleagues, which suggested that IL-4 induced by type 2 infections improves bystander-mediated protection by expanding the memory CD8 T cell pool^21^. Though their model used murine gammaherpesvirus (MuGHV) as a secondary, “bystander-activating” infection (as opposed to *L. monocytogenes* in this study), it too, requires IFN-γ for proper clearance^46,47^. These discordant results may be influenced by infection timing, wherein the increased duration between helminth and MuGHV infection may allow bystander CD8 T_mem_ to either become desensitized to IL-4 signals or to proliferate to a frequency great enough to compensate for impaired IFN-γ expression. Rolot, et al., observed helminth-enhanced IFN-γ levels and MuGHV clearance at 7 days post MuGHV infection when de novo pathogen-specific T cell responses peak, but not at the earlier timepoints when T cell responses are largely innate-like (days 1–3)^21^. Further, Rolot, et al., analyzed T cell functionality after stimulation with calcium ionophores, although bystander activation is independent of calcium flux^48^. Given this, we surmise that MuGHV-specific T cell responses may be misinterpreted as those driven by bystander activation. Nevertheless, additional studies will be necessary to address the durability of IL-4-mediated dysregulation of bystander responses and their intersection with antigen-specific T cell responses.

Our work demonstrates that IL-4 can augment bystander T cell activation, biasing cells for direct cytotoxicity. But cytokine expression varies across tissues: in human cadavers, the lung is a type 1 environment, and the skin is largely a type 2 (IL-4 enriched) environment^49,50^. Do these steady-state tissue environments curate bystander activation in an organ-specific manner? Lung-resident bystander T cells readily produce IFN-γ^15^, while those at the skin fail to express IFN-γ and instead kill targets directly^17,18^. While this may reflect the different pathogens used in these studies (extracellular bacteria and protozoans, respectively), bystander T cells in the skin remain biased for direct killing in alopecia models^22^. This suggests that tissue-intrinsic cytokine networks may dictate how bystander activation manifests—specifically elevated IL-4 signaling at the skin predisposing for direct cytotoxicity at the expense of IFN-γ secretion. Further research is needed to address if tissue-specific propensities for direct or indirect cytotoxicity are adaptations to better protect from the distinct pathogens that target these sites.

Our study is the first to identify mechanisms underlying strain-intrinsic differences in bystander activation—but these only manifest in SPF animals, as LCMV-experienced CD8 T_mem_ uniformly activated in response to IL-12/15/18. This extends our previous studies, which showed that IL-4 sensitivity affects CD8 T_mem_ differentiation in BALB/c mice but did not investigate bystander stimulation^51^. One could disregard the significance of these findings, given that SPF conditions are not observed in microbially-experienced human adults; but even antigen-experienced memory cells retain sensitivity to IL-4 that exceeds basal levels. Further, it is important to stress the SPF T cell repertoire mirrors human cord blood^52,53^. Given that neonates lack the antigen experience necessary for adaptive T_mem_ responses, these innate-like bystander T_mem_ responses are all the more critical for pathogen control in early life^54^. Therefore, genetic regulators of bystander activation may be important targets to enhance neonatal/postnatal immunity. Conversely, the fact that antigen experience overrides these strain-intrinsic determinants of bystander activation suggests studies using SPF animals may underestimate the contributions of bystander-activated CD8 T_mem_ to an immune response.

While basal IL-4 signaling reduces bystander activation in SPF BALB/c mice, *Il4ra* deletion does not elevate bystander activation in SPF BALB/c to levels observed in other strains. This suggests additional factors also control bystander activation. Nevertheless, formally dissecting the contributions of these signals in regulating bystander-mediated killing in vivo remains elusive, given the intractability of specifically depleting NK cells in the BALB/c model. Other animal models may prove useful in this regard; for instance, Lund and colleagues showed that recombinant-inbred “collaborative cross” animals display variability in their responsiveness to bystander-activating cytokines, albeit for unknown reasons^20^. Employing these tools alongside those that provide normal microbial experience will likely be of increasing importance to fully identify how bystander activation contributes to health and disease, as well as the therapeutic manipulation of these cells.

Overall, we demonstrate that protective responses following CD8 T_mem_ bystander activation can be impaired by steady-state, helminth-induced, and therapeutic IL-4 signals. We also highlight that propensities for bystander activation are determined by a combination of factors, including antigen experience and strain. Together, these underscore important considerations in manipulating bystander T cell responses therapeutically.

## Methods

### Mice

All animals were maintained in specific pathogen–free facilities at the University of Minnesota and infected in modified pathogen–free facilities. Experimental groups were nonblinded; animals were randomly assigned to experimental groups; and no specific method was used to calculate sample sizes.

We purchased all mice (C57BL/6J, CB6F1/J, BALB/cJ, C57BL/6J-*Ptprc^em6Lutzy^*/J [JAXBOY], and C57BL/6-Tg(*TcraTcrb*)1100Mjb/J [OT-I TCR transgenic]) from The Jackson Laboratory. We maintained congenically-distinct (i.e., CD45.1^+^) OT-I TCR transgenic mice by crossing C57BL/6-Tg(*TcraTcrb*)1100Mjb/J and C57BL/6J-*Ptprc^em6Lutzy^*/J mice. We euthanized mice in accordance with institutional protocols and subsequently collected spleens and LNs for experimentation.

### Pathogens and infections

We used an OVA-expressing VSV construct, administering 1 x 10^6^ PFU per mouse intravenously. We used wildtype *L. monocytogenes*. We grew *L. monocytogenes* in TSB media (+50µg/mL streptomycin) to the log phase of growth (determined by OD600), administering 1 x 10^3^ CFU per mouse intravenously. We propagated and prepared infective *H. polygyrus bakeri* third-stage larvae (L3) as previously described^45^, administering 200 L3/mouse via gavage. All pathogens were administered in sterile 1x PBS.

### IL-4 complexes

We complexed rIL-4 (Shenandoah Biotechnologies) with anti-IL-4 (clone: 11B11; BioXCell) at a 1:5 ratio by mass. We incubated complexes for 2–10 minutes at room temperature before diluting in 1x PBS for injections. We delivered 30µg IL-4c (5µg rIL-4, 25µg anti-IL-4) per mouse in a 200µL volume intraperitoneally.

### Developing OT-I memory mice

We prepared a single-cell suspension of spleen and lymph node cells that were harvested from female OT-I mice by mechanically passing tissue through a 70µm strainer. To enrich transgenic T cells, we used MACS with a CD8 Negative Selection Kit (Miltenyi Biotec). We adoptively transferred 1 x 10^4^ OT-I T cells in sterile 1x PBS i.v. per female C57BL/6 recipient and subsequently i.v. infected recipients with 1 x 10^6^ PFU VSV-OVA. We allowed OT-I memory T cells to contract to a stable memory pool (≥60d) before assaying tissues.

### T cell isolation and in vitro stimulation

We harvested spleen and LN from mice and mechanically prepared single-cell suspensions. To enrich bulk CD8 T cells from single-cell suspensions, we used mouse-specific CD8 T cell negative isolation “MojoSort” kits (BioLegend). We plated 0.5 to 1 x 10^6^ CD8 T cells per well in 96-well V-bottom tissue culture plates. We cultured cells in RP10 media (RPMI 1640 supplemented with 10% FBS, 2 mM L-glutamine, 100 U/mL penicillin-streptomycin, 1 mM sodium pyruvate, 0.05 mM β-mercaptoethanol, and 1 mM HEPES). We cultured T cells in RP10 with media alone, rIL-4 (Peprotech), rIL-7 (Peprotech), rIL-12p70 (Peprotech), rIL-15 (BioLegend), and/or rIL-18 (BioLegend) (each at 100ng/mL). For TCR stimulations, we coated plates with 10ug/mL, ea. of anti-CD3 (clone: 145-2C11) and -CD28 (clone: 37.51) (BioXCell) overnight at 4°C before decanting solution and plating CD8 T cells. We cultured cells at 37°C, 5% CO_2_, sampling cells at 24, 48, and 72 hours for flow staining. For intracellular cytokine staining, we added GolgiPlug (BD Biosciences) at a 1:1000 dilution 4 hours prior to cell harvest.

### In vivo bystander activation and ex vivo analysis

We i.v. infected mice with 1 x 10^3^ CFU wildtype *L. monocytogenes*. We infected mice at day 6 of *H. polygyrus* infection or immediately after IL-4c administration. At 24 and 72h timepoints, we i.v. injected mice with 3µg of allophycocyanin-conjugated anti-CD8b.2 (clone: 53-5.8) 3 minutes prior to euthanasia. We harvested blood, lymph nodes, and spleens and mechanically prepared single-cell suspensions by passing tissues through a 70µm filter. To remove contaminating red blood cells, we incubated blood and spleen suspensions in ACK buffer for 5 minutes, and then proceeded to conduct staining for flow cytometric analysis.

### Flow cytometry

For in vitro and ex vivo analyses we conducted all stains at 4°C. All reagent information is included in **Supplemental table 1**. We conducted LIVE/DEAD fixable blue viability dye (Thermo Fisher) staining in 1x PBS. For surface staining, we utilized brilliant staining buffer (BD Biosciences) as the stain diluent. Surface stains were for 25 minutes and were extended to 60 minutes for tetramer staining. For intracellular cytokine staining, we fixed cells in 1x FOXP3/Transcription Factor Fixation/Permeabilization buffer (Cytek) and conducted intracellular/nuclear stains using 1x Flow Cytometry Perm Buffer (Cytek) as the diluent. For panels interrogating APCs, we fixed cells in Cytofix/Cytoperm (BD Biosciences) and conducted intracellular staining using 1x Perm/Wash (BD) as the diluent. Fixation was always for 20 minutes and intracellular staining for 30 minutes. We resuspended cells in FACSWash (1x PBS, 2% FBS, 0.2% sodium azide) and acquired events on a LSRFortessa (BD Biosciences), which we analyzed using FlowJo v10 (BD Biosciences). We conducted statistical testing using Prism v10 (GraphPad).

## Acknowledgements

We thank Dr. Joseph Urban for providing *H.polygyrus* larvae and Drs. Sara Hamilton-Hart and Courtney Matson for critical review of the manuscript. N.N.J. is a Damon Runyon Fellow supported by the Damon Runyon Cancer Research Foundation (Grant No. DRG-2427-21). N.J.M. is an NCI K00 Fellow supported by the National Cancer Institute (Grant No. K00 CA245735). This work was supported by R01AI38903 (S.C.J.). Monomer and tetramer reagents were obtained through the NIH Tetramer Core Facility.

