## Supplemental figures and tables for "Steady-state, therapeutic, and helminth-induced IL-4 compromise protective CD8 T cell bystander activation"

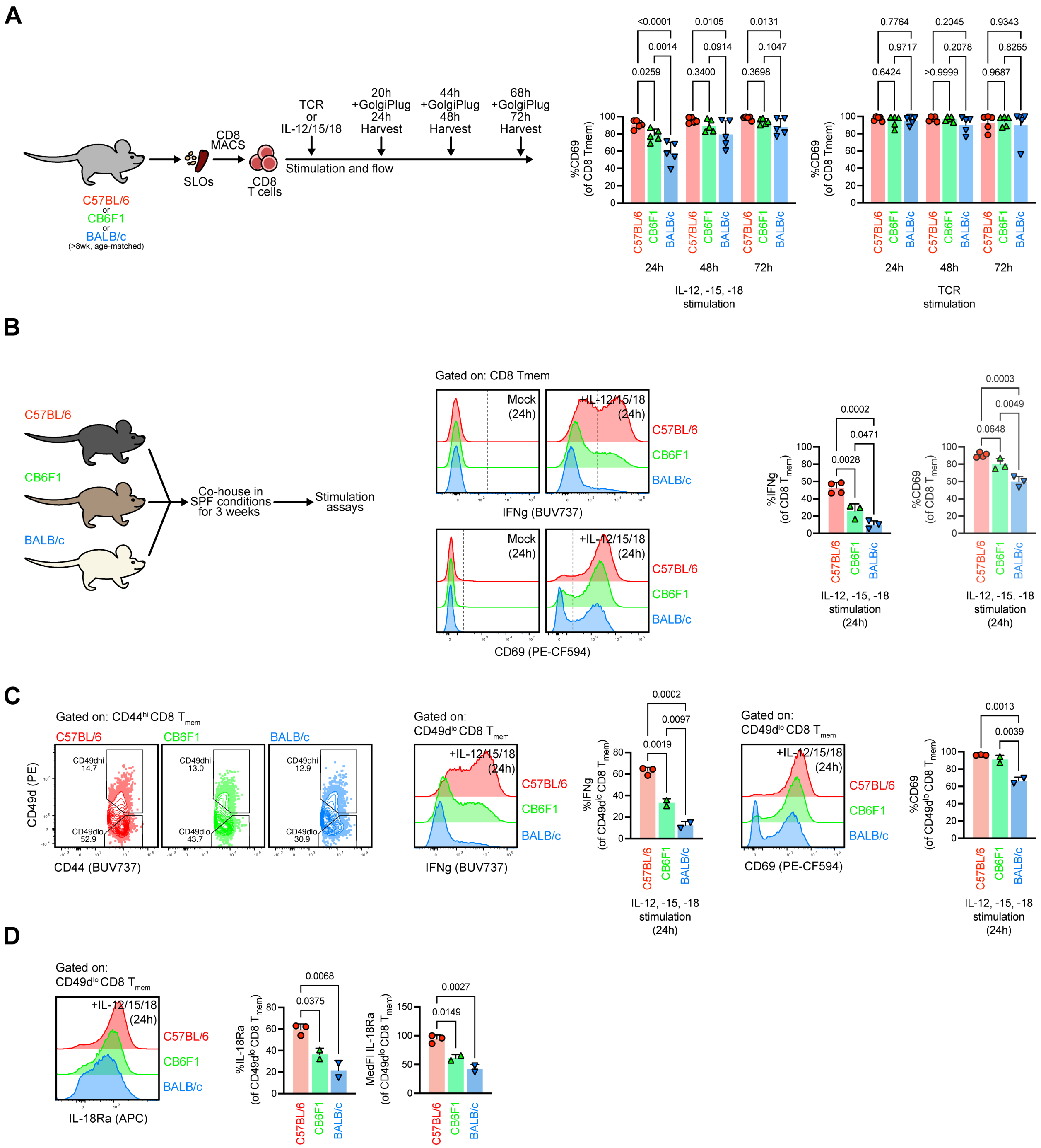


### Supplementary figure 1. Bystander activating cytokines differentially activate CD8 $T_{mem}$ from SPF laboratory mice.

**A** CD69 expression in memory phenotype ($CD44^{hi}$) CD8 T cells after stimulation with bystander-activating cytokines (IL-12/15/18, 100ng/mL, ea.) or TCR agonists. **B** Expression of IFN-γ and CD69 in CD8 $T_{mem}$ in CD8 $T_{mem}$ from SPF co-housed mice. **C** Representative gating of $CD49d^{lo}$

virtual memory CD8 T cells ($T_{VM}$) and expression of IFN-$\gamma$ and CD69 after exposure to IL-12/15/18. **D** Expression of IL-18Ra in CD8 $T_{mem}$ from SPF co-housed animals. Points in **A–D** depict unique animals (n=2–5) animals across (**A** and **B**) 2 and (**C** and **D**) 1 technical replicate.

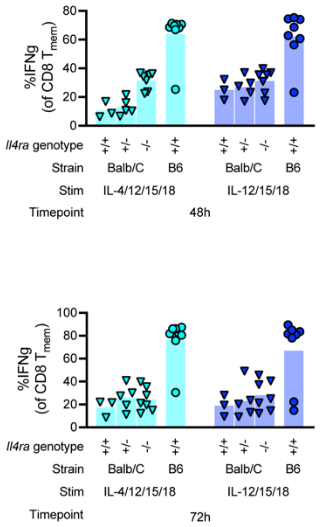


**Supplementary figure 2. Disrupting steady-state IL-4 signaling in BALB/c CD8 $T_{mem}$ partially restores response to bystander-activating cytokines**

We stimulated and analyzed cells from BALB/c *Il4ra* variants as outlined in **Figure 2B**.

Expression of IFN-$\gamma$ in CD8 $T_{mem}$ at 72h and 120h timepoints of stimulation with IL-4/12/15/18 or IL-12/15/18 (100ng/mL, ea.). Each symbol in  depicts cells from an individual animal in a unique stimulation condition (n=3–8 per strain/geneotype) across 2 technical replicates.

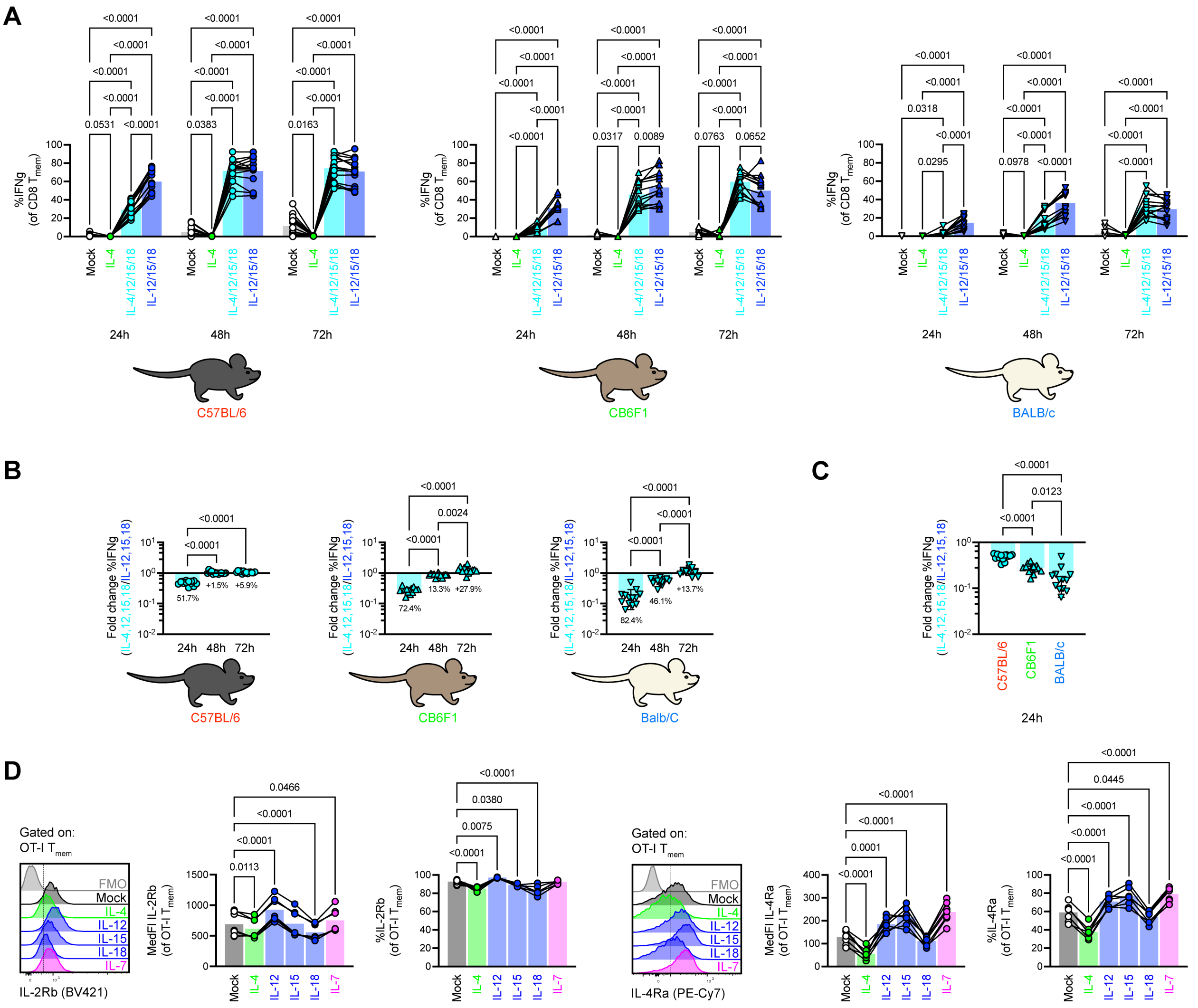


**Supplementary figure 3. IL-4 limits IL-18Ra expression and IFN-γ expression elicited by bystander-activating cytokines.**

**A.** We stimulated cells from SPF C57BL/6, CB6F1, and BALB/c mice as outlined in **Fig. 2A** with bystander activating cytokines (IL-12/15/18, 100ng/mL, ea.) in the presence or absence of IL-4 (100ng/mL) and interrogated cells using flow. Expression of IFN-γ in CD8 $T_{mem}$ across SPF strains. **B.** IL-4-mediated fold reduction of IFN-γ elicited by bystander-activating cytokines. **C.** Cross-strain comparison of IL-4-mediated fold reductions of IFN-γ during in vitro bystander activation. **D** and **E.** We stimulated cells as outlined in **Fig. 2C** and interrogated cells using flow. Expression of **D** IL-2Rb and **E** IL-4Ra in OT-I $T_{mem}$.

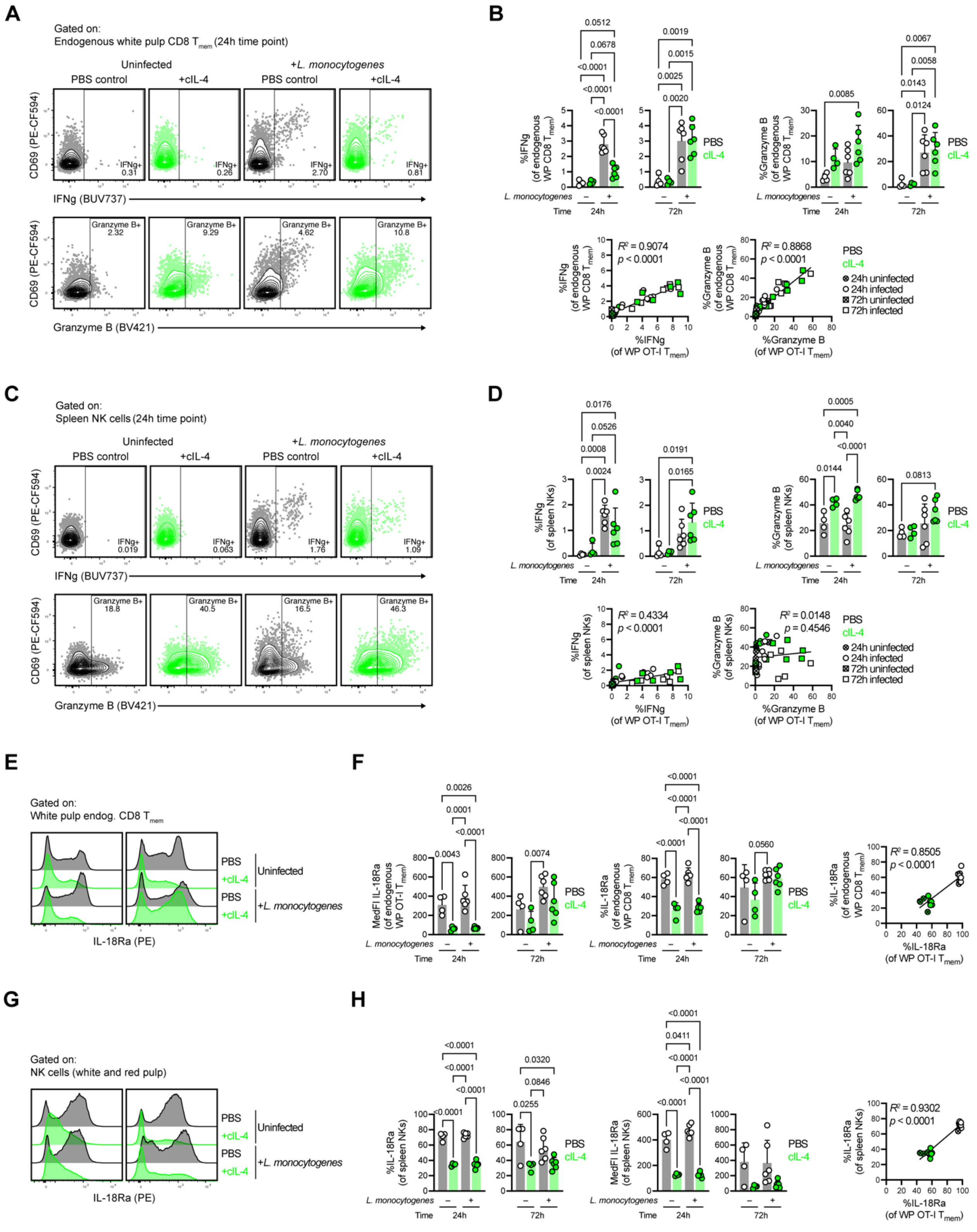

**Supplementary figure 4. IL-4 complex therapies impair protective functions of endogenous CD8 $T_{mem}$ and NK cells.**

**A–H** We treated mice with IL-4c and infected with *L. monocytogenes* as described in **F4A. A, B** IFN-γ and granzyme B expression in endogenous CD8 $T_{mem}$ at the splenic white pulp (WP). **C** Correlation of IFN-γ and granzyme B levels in WP OT-I $T_{mem}$ and WP endogenous CD8 $T_{mem}$. **D, E** IFN-γ and granzyme B expression in bulk splenic NK cells. **F** Correlation of IFN-γ and granzyme B levels in WP OT-I $T_{mem}$ and splenic NK cells. **G** IL-18Ra expression in WP endogenous CD8 $T_{mem}$. **H** Correlation of IL-18Ra expression in WP OT-I $T_{mem}$ and WP endogenous CD8 $T_{mem}$. **I** IL-18Ra expression in splenic NK cells. **J** Correlation of IL-18Ra expression in WP OT-I $T_{mem}$ and splenic NK cells. Each point in **B**, **D**, **F**, and **H** represent an individual animal (n=3–6 per timepoint and condition) across 2 technical replicates. Indicated statistical significance was calculated by ordinary one-way ANOVA with Tukey's multiple comparisons test or linear regression.

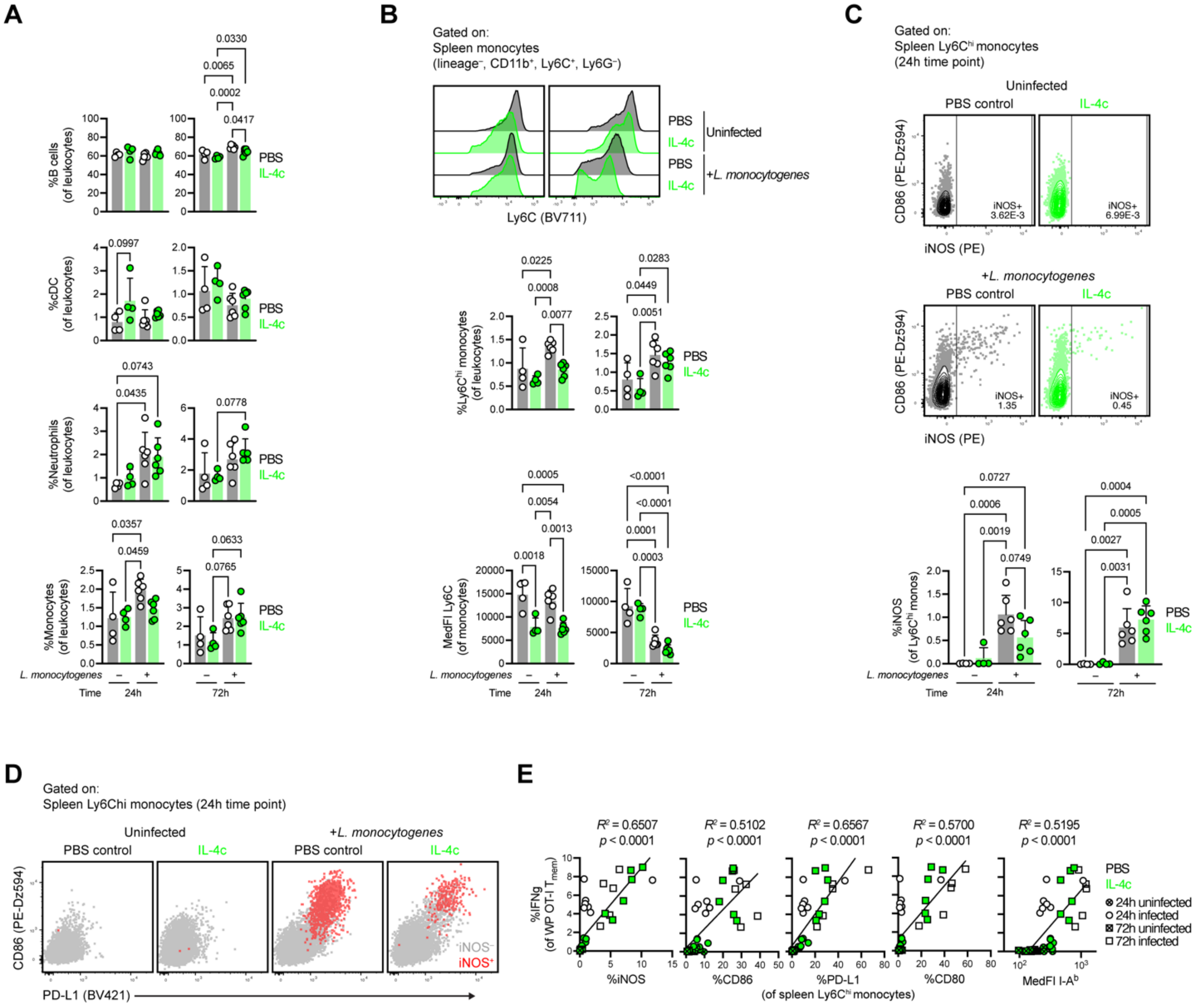


**Supplementary figure 5. IL-4c impairs IFN-γ–mediated monocyte instruction**

We treated mice with IL-4c and infected with *L. monocytogenes* as described in **F4A. A** Frequencies of splenic antigen-presenting cell (APC) subsets during IL-4c treatment and/or *L. monocytogenes* infection. **B** Ly-6C expression in splenic monocytes during IL-4c treatment and/or *L. monocytogenes* infection. **C** iNOS expression in Ly-6C$^{hi}$ splenic monocytes during IL-4c treatment and/or *L. monocytogenes* infection. **D** Representative plots of IFN-γ–induced gene expression in iNOS$^{+}$ (red) and iNOS$^{-}$ (grey) splenic Ly-6C$^{hi}$ monocytes. **E** Correlation of IFN-γ–induced gene expression in splenic Ly-6C$^{hi}$ monocytes with IFN-γ expression in splenic WP OT-I T$_{mem}$. Each point in **A**, **B**, **C**, and **E** represent an individual animal (n=3–6 per timepoint and condition) across 2 technical replicates. Indicated statistical significance was calculated by ordinary one-way ANOVA with Tukey’s multiple comparisons test or linear regression.

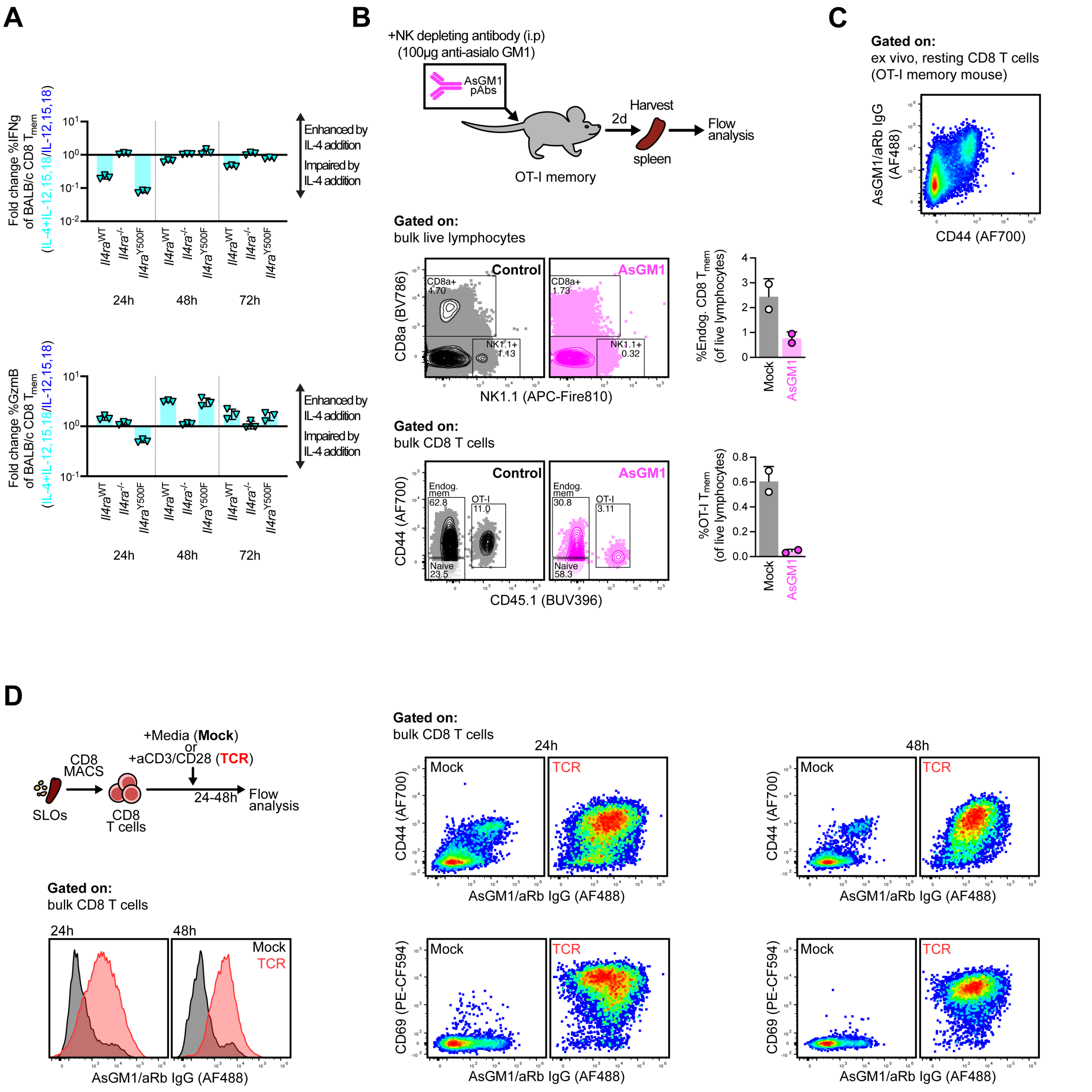


**Supplementary figure 6. IL-4 augments bystander activation of CD8 $T_{mem}$ in a PI3-K-independent manner.**

**A** We stimulated CD8 T cells isolated from wildtype, *Il4ra*$^{-/-}$, and *Il4ra*$^{Y500F}$ BALB/c mice and stimulated with IL-4 (100ng/mL) and/or IL-12/15/18 (100ng/mL) in a setup similar to **Figure 5A** and interrogated cells using flow. IL-4-mediated fold change of IFN-γ (top) and granzyme B (bottom) in CD8 $T_{mem}$ stimulated with IL-12/15/18. **B** We treated animals with depleting AsGM1 antibodies and measured NK and CD8 $T_{mem}$ populations. **C** Representative plot depicting

AsGM1 expression in bulk CD8 T cells from an OT-I memory mouse at homeostasis. **D** We isolated CD8 T cells from SPF animals and stimulated with media or TCR agonists and measured AsGM1 expression. Representative plots depict AsGM1 expression in bulk CD8 T cells. Each symbol in **A** depicts cells from an individual animal in a unique stimulation condition (n=3) across 1 technical replicate. Each symbol in **B** depicts a single animal (n=2 per condition) in 1 technical replicate. Plots in **C** and **D** are representative of 3 animals across 1 technical replicate.

**Supplementary table 1. Flow cytometry reagents**

| Reagent | Conjugate | Clone | Vendor | Dilution |
|---|---|---|---|---|
| **In vivo labeling reagents** | | | | |
| CD8b.2 | APC | | ThermoFisher | 3µg/mouse |
| **Viability stain reagents** | | | | |
| Blue viability dye (BViD) | UV450 | N/A | ThermoFisher | 1:500 |
| **Surface stain reagents** | | | | |
| TruStain FcX | NA | 93 | BioLegend | 1:200 |
| CD132 (yc) | Biotin | TUGm2 | ThermoFisher | 1:200 |
| PD-L1 (CD274) | Biotin | 10F.9G2 | BioLegend | 1:100 |
| CD45.1 | BUV395 | A20 | BD Biosciences | 1:200 |
| CD62L | BUV395 | MEL-14 | BD Biosciences | 1:200 |
| CD80 | BUV395 | 16-10A1 | BD Biosciences | 1:100 |
| CD19 | BUV737 | 1D3 | BD Biosciences | 1:200 |
| Streptavidin | BV421 | NA | BD Biosciences | 1:500 |
| CD122 (IL-2Rb) | BV421 | TM-β1 | BD Biosciences | 1:200 |
| ICOS | BV421 | C398.4A | BioLegend | 1:200 |
| CD44 | BV510 | IM7 | BD Biosciences | 1:500 |
| F4/80 | BV510 | BM8 | BioLegend | 1:500 |
| NKp46 | BV510 | 29A1.4 | BioLegend | 1:200 |
| PD-1 (CD279) | BV605 | 29F.1A12 | BioLegend | 1:200 |
| Ly-6G | BV605 | 1A8 | BD Biosciences | 1:200 |
| NKG2D (CD314) | BV711 | CX5 | BD Biosciences | 1:200 |
| Ly-6C | BV711 | HK1.4 | BioLegend | 1:500 |
| CD8a | BV786 | 53-6.7 | BD Biosciences | 1:200 |
| CD11c | BV786 | N418 | BioLegend | 1:500 |
| KLRG1 | FITC | 2F1 | Cytek | 1:200 |
| NK1.1 | FITC | PK136 | BioLegend | 1:100 |
| I-A$^{b}$ | FITC | AF6-120.1 | BD Biosciences | 1:100 |
| D$^{b}$ LCMV GP33-41 | AF488 | NA | NIH Tetramer Core | 1:100 |
| L$^{d}$ LCMV NP118-126 | AF488 | NA | NIH Tetramer Core | 1:100 |
| TCRb | PerCP-Cy5.5 | H57-597 | BioLegend | 1:200 |
| CD49d | PE | 9C10 | BioLegend | 1:200 |
| 4-1BB (CD137) | PE | 17B5 | BioLegend | 1:100 |
| CD218a (IL-18Ra) | PE | A17071D | BioLegend | 1:100 |
| D$^{b}$ LCMV NP396-404 | PE | NA | NIH Tetramer Core | 1:200 |
| Streptavidin | PE-Dz594 | NA | BioLegend | 1:500 |
| CD69 | PE-CF594 | H1.2F3 | BD Biosciences | 1:200 |
| CD86 | PE-Dz594 | GL-1 | BioLegend | 1:100 |
| CD19 | PE-Cy5.5 | eBio1D3 | ThermoFisher | 1:200 |
| CD124 (IL-4Ra) | PE-Cy7 | I015F8 | BioLegend | 1:100 |
| D$^{b}$ LCMV GP33-41 | APC | NA | NIH Tetramer Core | 1:200 |
| CD218a (IL-18Ra) | AF647 | A17071D | BioLegend | 1:100 |
| ICOS | AF647 | C398.4A | BioLegend | 1:200 |
| CD11b | AF700 | M1/70 | BD Biosciences | 1:500 |
| CD44 | AF700 | IM7 | BD Biosciences | 1:200 |
| CD8a | APC-e780 | 53-6.7 | ThermoFisher | 1:200 |
| CD45.2 | APC-e780 | 104 | ThermoFisher | 1:200 |
| NK1.1 | APC-fire810 | S17016D | BioLegend | 1:100 |
| **Intracellular stain reagents** | | | | |
| IFN-γ | BUV737 | XMG1.2 | BD Biosciences | 1:200 |
| Granzyme B | BV421 | GB11 | BD Biosciences | 1:200 |
| Granzyme B | PE | GB11 | ThermoFisher | 1:200 |
| iNOS | PE | REA982 | Miltenyi Biotec | 1:200 |
| Ki-67 | PE-Cy7 | 16A8 | BioLegend | 1:500 |
| Granzyme B | APC | GB11 | ThermoFisher | 1:200 |
